# The Ins and Outs of Manganese: ZIP14 facilitates the efflux of excess manganese from the brain

**DOI:** 10.1101/2025.07.02.662898

**Authors:** J Zou, TL Thorn, Z Wang, Y Wang, TB Aydemir

**Affiliations:** Cornell University, Division of Nutritional Sciences, 244 Garden Avenue 14853 Ithaca NY

**Author notes:** **Correspondence:** Tolunay Beker Aydemir, Cornell University, Division of Nutritional Sciences, 244 Garden Avenue 14853 Ithaca NY, 4046446919.

**Keywords:** Blood-brain barrier, metal transporter, basolateral localization, metal homeostasis, SLC39A14

## Abstract

Manganese (Mn) is essential for many enzymatic processes in the brain; however, Mn overload can lead to neurotoxicity and behavioral deficits. The blood-brain barrier (BBB), comprised of polarized endothelial cells, tightly regulates metals in and out of the brain. ZIP14 (*SLC39A14*) is a metal transporter more recently found to transport Mn, with mutations in human *SLC39A14* resulting in brain Mn accumulation and neurological deficits. However, ZIP14’s precise localization and role in BBB endothelial cells remain unclear. Here, we show *in vivo* ZIP14 expression in BBB endothelial cells, which upregulates following Mn supplementation. Using expansion microscopy, we observed a shift in ZIP14 localization from an equal apical-basolateral distribution to predominantly basolateral after Mn exposure. Endothelial-specific *Zip14* KO (EKO) mice exhibited impaired Mn efflux from the brain and increased brain Mn accumulation after nasal delivery and dietary Mn supplementation. *In vitro* studies using primary endothelial cells from EKO mice and ZIP14-overexpressing hCMEC/D3 cells confirmed that ZIP14 primarily mediates basolateral-to-apical Mn transport. Collectively, our results demonstrate that ZIP14 is critical for brain Mn clearance, highlighting its potential as a therapeutic target to mitigate Mn-induced neurotoxicity.

**Significance Statement:** Our study reveals that the endothelial metal transporter ZIP14 plays a critical role in removing excess Mn from the brain. Using expansion microscopy, genetic manipulation, metallomics, and transport studies employing radiolabeled Mn, we demonstrate that ZIP14 undergoes strategic relocalization in response to Mn exposure and primarily functions in basolateral-to-apical transport. Deletion of endothelial ZIP14 leads to brain Mn accumulation, establishing its fundamental role in protecting against Mn-induced neurotoxicity. This finding provides mechanistic insight into how the brain protects itself against Mn overload and why ZIP14 mutations in humans result in brain Mn accumulation.

## Introduction

Manganese (Mn) is an essential micronutrient and trace mineral required for various physiological processes, including amino acid metabolism, the urea cycle, carbohydrate metabolism, glycosylation, and antioxidant activity (1). As a cofactor for many metalloproteins, Mn plays a crucial role in maintaining cellular function (2). However, in excess, Mn can be neurotoxic. In humans, overexposure of Mn can result in parkinsonism, presenting with cognitive and motor symptoms similar to Parkinson’s Disease, such as tremors, muscle weakness, and worsened memory and attention (3–15).

Mn overexposure is on the rise due to increased exposure from occupational and lifestyle hazards, like toxic fume inhalation, contaminated water near industrial regions, and intravenous drug use (16). In addition to environmental exposures, mutations in the solute carrier (SLC) family of metal transporters have also been shown to lead to Mn accumulation. In humans, *SLC39A14* mutations can lead to childhood-onset parkinsonism with hypermanganesemia, presented as dystonia and later as tremor and bradykinesia (17–26). Mutations in *SLC39A14* have been associated with Mn toxicity in both humans and animal models (27–32). Similarly, mutations in *SLC30A10*, which encodes ZnT10, can also result in Mn overload, manifesting in childhood or adulthood (33–37). Since both ZIP14 and ZnT10 are expressed in liver and intestines and facilitate Mn excretion, the absence of these will result in systemic accumulation of Mn.

While it is well established that excess Mn targets the brain, the transport dynamics and transporters involved in Mn homeostasis are not fully understood. The transporter of interest in this study is ZIP14. Initially thought to primarily transport zinc, ZIP14 is expressed on the plasma membrane and organelles, and transports divalent metals, including Mn, from the extracellular or organelle spaces into the cell cytosol (17, 38–40). To investigate the contributions of ZIP14 in brain accumulation, we and other researchers have used whole-body *Zip14* knockout (KO) mice as a model of *SLC39A14* loss-of-function mutations, which led to significant brain Mn accumulation (17, 27–32). These findings suggested that ZIP14 may not be crucial to Mn uptake into the brain. Our subsequent studies with intestine-specific *Zip14* KO mice revealed significant Mn accumulation in the blood and brain (41) while liver-specific *Zip14 KO* did not cause systemic or brain Mn accumulation (32), indicating that intestinal Mn excretion is critical for the maintenance of Mn homeostasis. These findings were supported by liver and intestine double KO studies (42). However, no mouse model has directly studied ZIP14’s role in Mn transport directionality at the BBB *in vivo*. To address this gap, we generated an endothelial-specific *Zip14* KO mouse model to investigate the role of BBB ZIP14 in brain Mn accumulation. With this mouse model, we assessed ZIP14’s role in the directional transport of Mn by administering a radioactive Mn tracer either directly to the brain via the nasal route to determine efflux or subcutaneously to determine uptake. We also performed radioactive tracer studies in primary endothelial cells isolated from EKO mouse brains and hCMEC/D3 cell line to further explore the directionality of Mn transport by ZIP14.

## Results

### ZIP14 is expressed in brain endothelial cells

We first examined ZIP14 expression in brain endothelial cells *in vivo* using immunofluorescence. Colocalization of ZIP14 staining with PECAM1, an endothelial cell marker, demonstrated ZIP14 expression in brain endothelial cells. This was verified in four complementary ways: *in vivo* whole brain sections, *ex vivo* isolated brain microvessels, and primary brain endothelial cells cultured in both 2D monolayers and 3D tubules formed on Matrigel. In the whole brain sections, we observed colocalization of ZIP14 and PECAM1 in both longitudinal and transverse vessel sections (Figure 1A). In isolated microvessels, ZIP14 and PECAM1 colocalized, and we also noted regions of ZIP14 staining without PECAM1, likely corresponding to remnants of astrocyte endfeet that wrap around the BBB endothelial cells (Figure 1B) (43, 44). We generated a 3D reconstruction from z-stacks of a brain vessel, acquired with expansion microscopy, from fl/fl mice following Mn supplementation. Similarly, ZIP14 expression was detected throughout the vessel, appearing as discontinuous and punctate rather than a continuous pattern (*Supplementary Figure 2B*).

**Figure 1.**
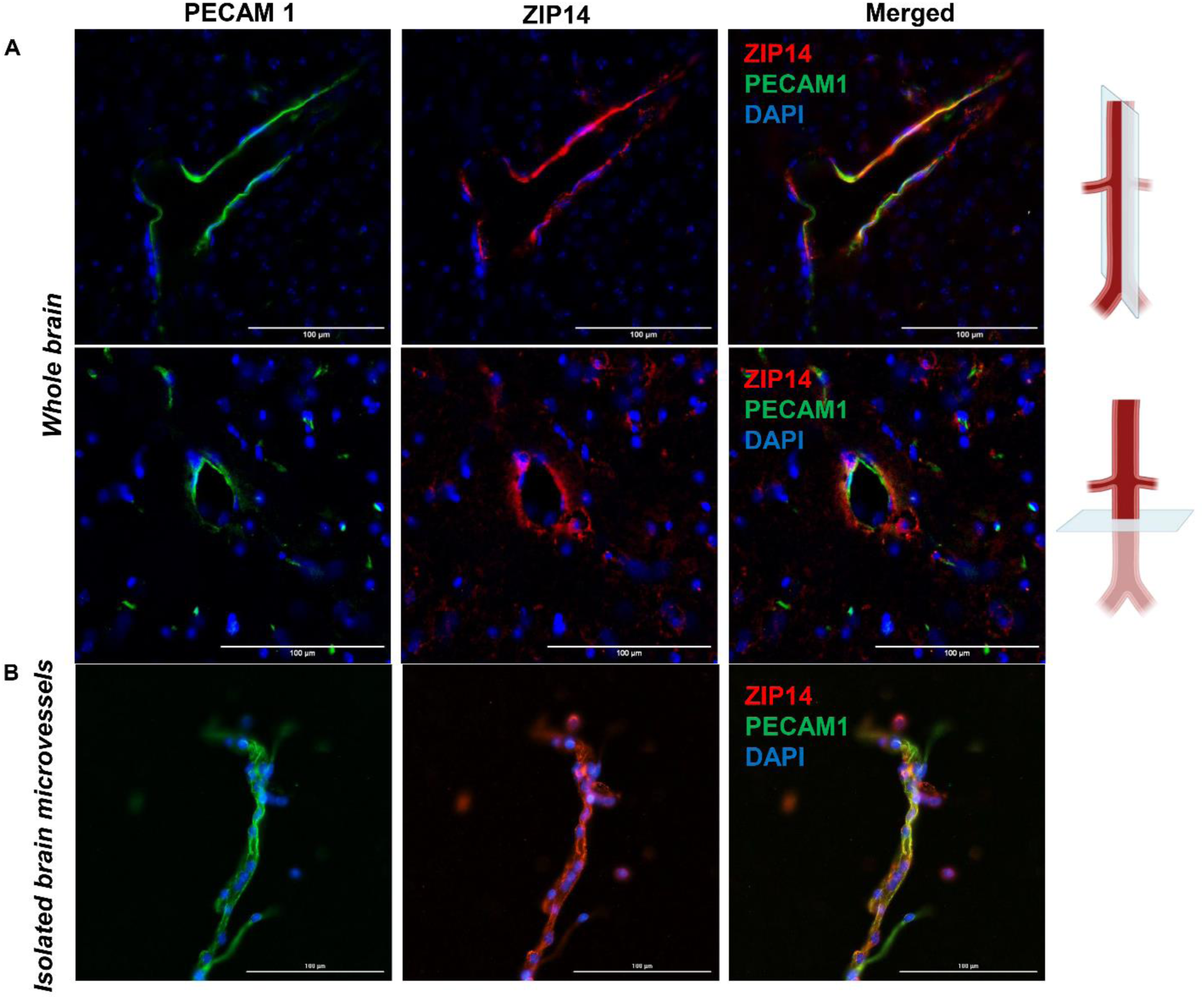
In vivo ZIP14 expression in brain endothelial cells. Representative immunofluorescent images showing ZIP14 (red) co-localized with the endothelial marker, PECAM1 (green) in A) whole mouse brain sections cut longitudinally (top) and transversely (bottom); B) isolated mouse brain microvessels. Scale bars indicate image magnification.

Primary endothelial cells were isolated from mouse brains (Figure 2A) and in the 2D culture of primary endothelial cells, ZIP14 was localized both on the plasma membrane and intracellular organelles (Figure 2B). When grown on Matrigel, these primary endothelial cells formed 3D tubules (Figure 2C). At high magnification, individual tubules formed by two endothelial cells can be visualized, showing clear colocalization of ZIP14 and PECAM1 (Figure 2C).

**Figure 2.**
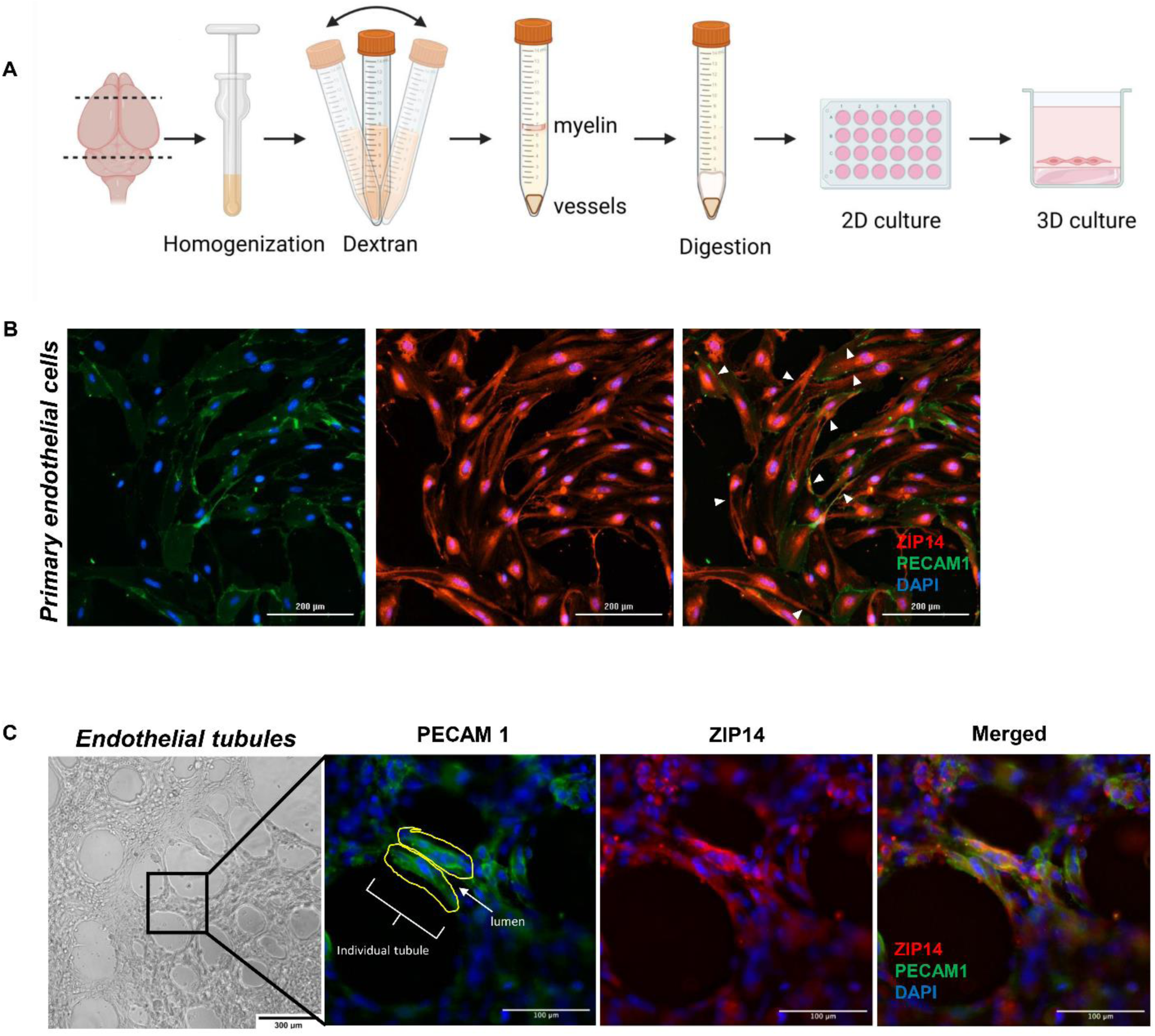
In vitro ZIP14 expression in primary brain endothelial cells. A) Schematic of isolation procedure for primary brain endothelial cells: mouse cerebrum was homogenized with a douncer, followed by myelin separation with 18% dextran and enzymatic digestion of the vessel pellet with collagenase/dispase. The resulting endothelial cell suspension was seeded onto 24-well plates. B) Representative immunofluorescent images show ZIP14 (red) co-localized with the endothelial marker, PECAM1 (green) in 2D-cultured primary endothelial cells. Arrowheads point to ZIP14 signal on the plasma membrane. C) Primary endothelial cells seeded on Matrigel formed a network of tubules, as shown in the brightfield image. Representative images display ZIP14 and PECAM1 staining in one individual tubule. The arrow indicates the lumen, and the yellow outline highlights endothelial cells forming the tubule. Scale bars indicate image magnification.

### ZIP14 expression increases in response to manganese

It is known that ZIP14 expression is dynamic and can change with Mn exposure, as shown in human hepatocytes (45) and zebrafish (17). However, this has not been fully explored in brain endothelial cells. We observed an increase in ZIP14 expression with Mn overexposure, as visually demonstrated through immunofluorescent staining in whole brain sections after subchronic nasal Mn supplementation (Figure 3A-B). The nasal Mn supplementation method was validated by a significant, more than two-fold increase in brain Mn levels (Figure 3E). Although there were also significant increases in Zn and Fe levels, the extent of increase was relatively small. The change in ZIP14 expression was replicated *in vitro* in primary brain endothelial cells and with a greater ZIP14 signal after MnCl_2_ supplementation (Figure 3C-D). Accompanying the increase in ZIP14 signal, we saw greater uptake of Mn into primary endothelial cells as extracellular Mn increased (Figure 3F). This is expected as ZIP14 on the plasma membrane serves as an importer of Mn into the cell.

**Figure 3.**
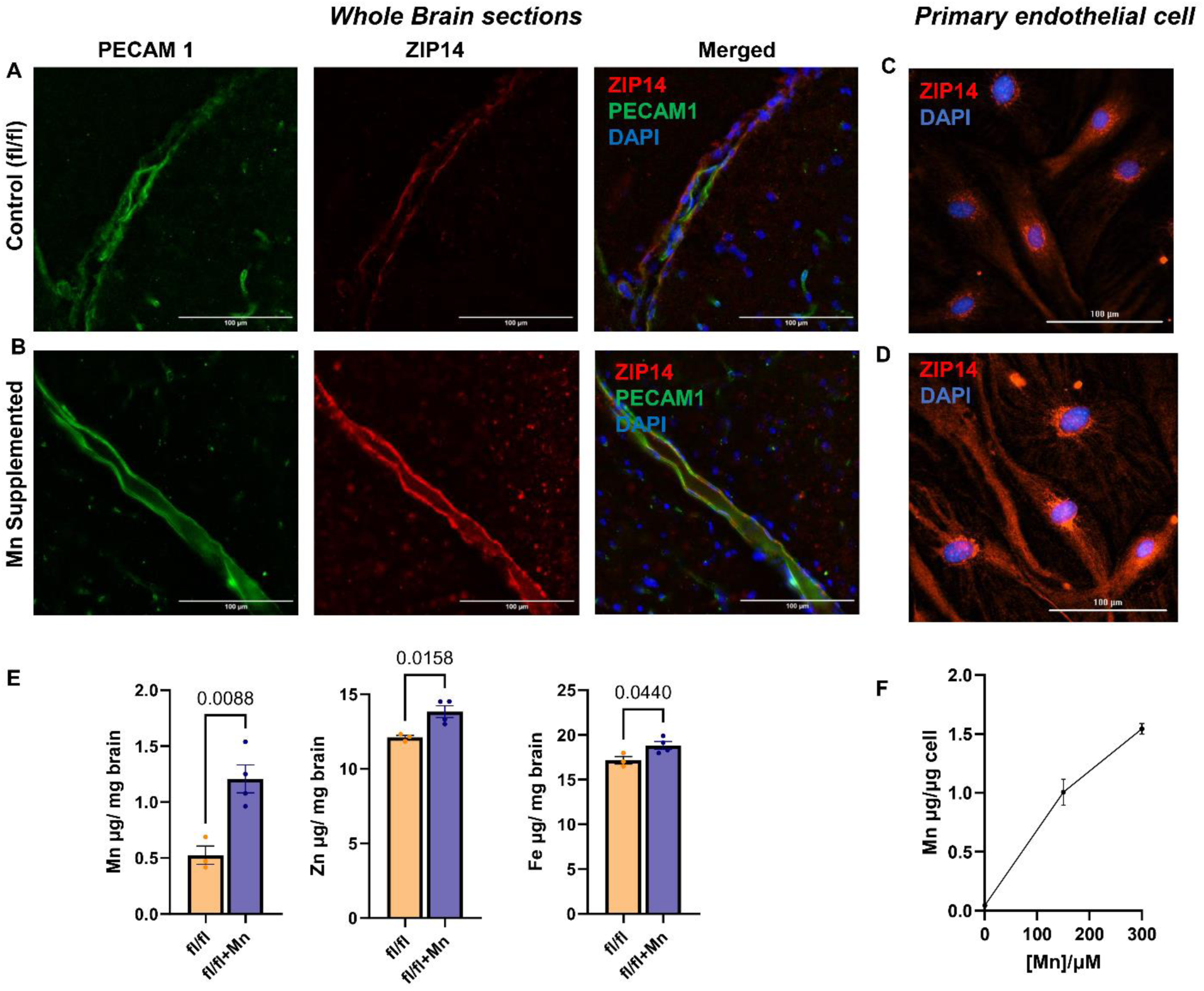
ZIP14 expression is upregulated in response to Mn exposure. Representative immunofluorescent images of ZIP14 (red) and endothelial marker PECAM1 (green) in whole brain sections from A) untreated fl/fl control mice and B) fl/fl mice after 4 weeks of nasal Mn supplementation. Representative immunofluorescent images of ZIP14 (red) in primary endothelial cells C) vehicle control or D) 300μM MnCl_2_ for 72hrs. E) Brain Mn, Zn, and Fe levels in fl/fl control mice (n=3) and fl/fl mice treated with 4 weeks of nasal Mn supplementation (n=4). F) Mn levels in primary endothelial cells treated with 0, 150, 300μM MnCl_2_ for 72hrs (n=3). Data presented as mean±SEM (Student’s t-test). Scale bars indicate image magnification.

### ZIP14 expression is enriched in the basolateral surfaces following Mn exposure

Given the polarized nature of brain endothelial cells, we sought to determine the localization of ZIP14, whether it resides on the apical or basolateral side, indicating Mn uptake or efflux, respectively. Here, we used expansion microscopy and Matrigel tubule formation to investigate *in vivo* and *ex vivo* distribution of ZIP14.

Using immunofluorescent staining and expansion microscopy, we identified cross-sections of blood vessels in brain sections from fl/fl mice with or without Mn supplementation and observed localization of ZIP14 in endothelial cells surrounding the lumen (Figure 4A-B). ZIP14 signal intensity was significantly higher in Mn-treated fl/fl mice compared to fl/fl control (Figure 4C). This is expected as we found *Zip14* mRNA expression was also significantly upregulated in brain microvessels isolated from Mn-treated fl/fl mice (Figure 4D). Endothelial cells making up the vessel had apical and basolateral surfaces (Figure 4E). Quantification of ZIP14 intensity in untreated vessels revealed approximately equal distribution of ZIP14 on the apical and basolateral sides of the vessels (Figure 4F; *Supplementary Figure 1*). However, following Mn supplementation, ZIP14 signal shifted significantly towards the basolateral side (Figure 4F; *Supplementary Figure 2A*).

**Figure 4.**
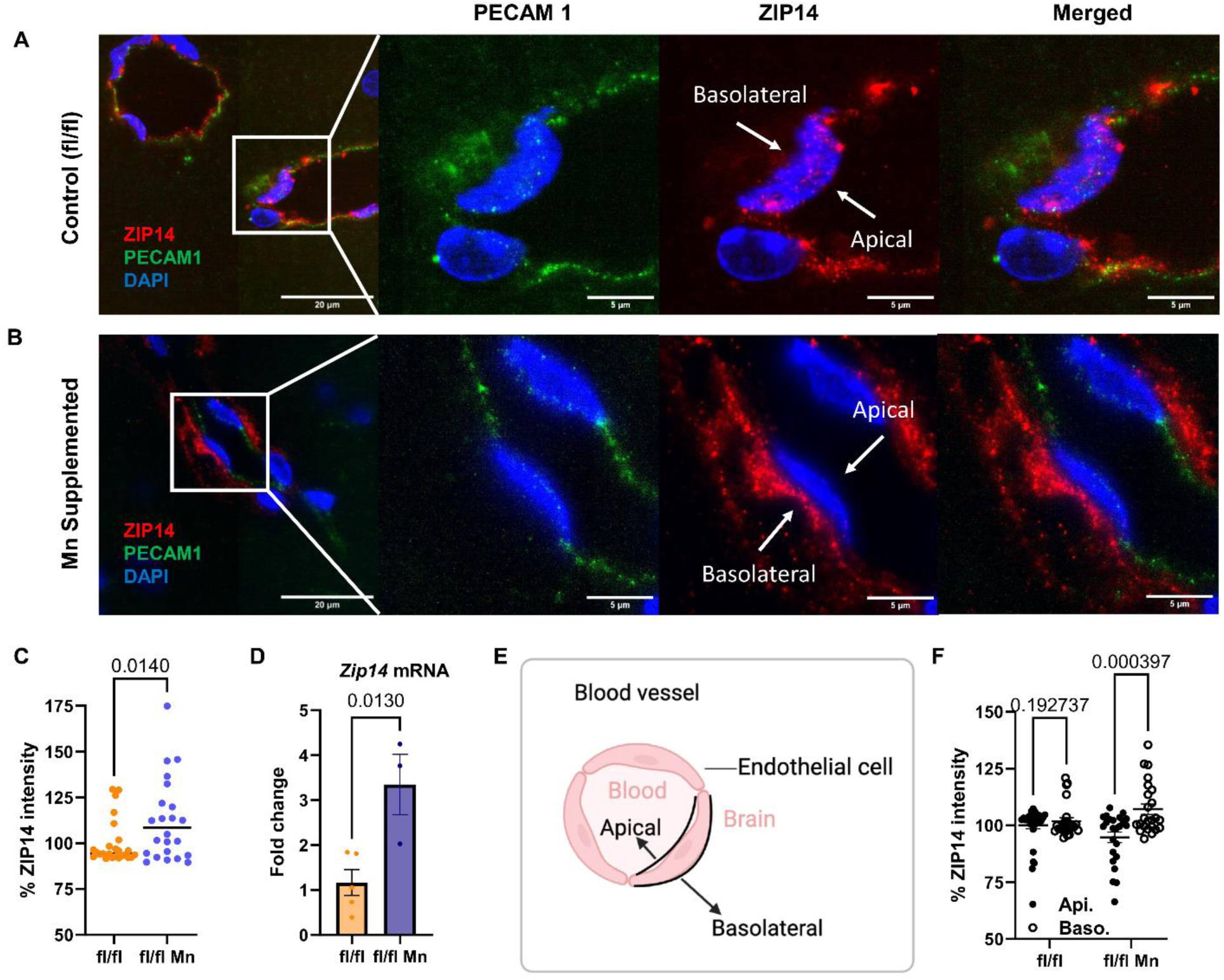
ZIP14 shifts to basolateral localization after Mn supplementation. Representative immunofluorescent images of ZIP14 (red) and endothelial marker PECAM1 (green) in expanded (3-fold) whole-brain sections from A) fl/fl control mice receiving nasal vehicle control (water) delivery and B) fl/fl mice supplemented with nasal Mn for 4 weeks. C) Quantification of total ZIP14 immunofluorescence intensity (combined apical and basolateral signals) in endothelial cells from fl/fl and Mn-supplemented brain vessels. Values are expressed relative to fl/fl (set to 100%). D) *Zip14* mRNA expression in brain microvessels isolated from fl/fl control mice (vehicle-treated; n=5) and Mn-supplemented fl/fl mice (n=3). E) Schematic showing the cross-section of a vessel and the apical and basolateral surfaces of surrounding endothelial cells. F) ZIP14 localization on apical versus basolateral surface of brain vessel endothelial cells was quantified in fl/fl and Mn supplemented fl/fl mice. Values are expressed relative to fl/fl apical values (set to 100%). Data represent mean±SEM (8 vessels/group, multiple endothelial cells analyzed per vessel; Student’s t-test). Scale bars indicate image magnification.

### Generation of endothelial-specific *Zip14* knockout mouse model

To specifically examine the role of ZIP14 in the BBB endothelial cells, we generated endothelial-specific *Zip14* knockout mice (EKO) that had *Zip14* deleted in *tie2*-specific cells. A Cre-lox system was used, as shown in the representative genotype PCR assay (Figure 5A). We confirmed the knockout through western blot analysis of isolated brain microvessels from EKO mice (Figure 5B). Furthermore, we observed a significant reduction in *Zip14* mRNA levels in EKO primary endothelial cells, with no change in intestinal epithelial cells, which are known to highly express ZIP14 (Figure 5C). Immunofluorescent staining further validated a reduction in ZIP14 signal in brain microvessels from EKO mice (Figure 5D). These results confirm that *Zip14* was knocked out in EKO mice. Notably, ZIP14 was not completely absent in protein or mRNA expression, likely due to its expression in astrocyte endfeet that are present in blood vessels of the brain.

**Figure 5.**
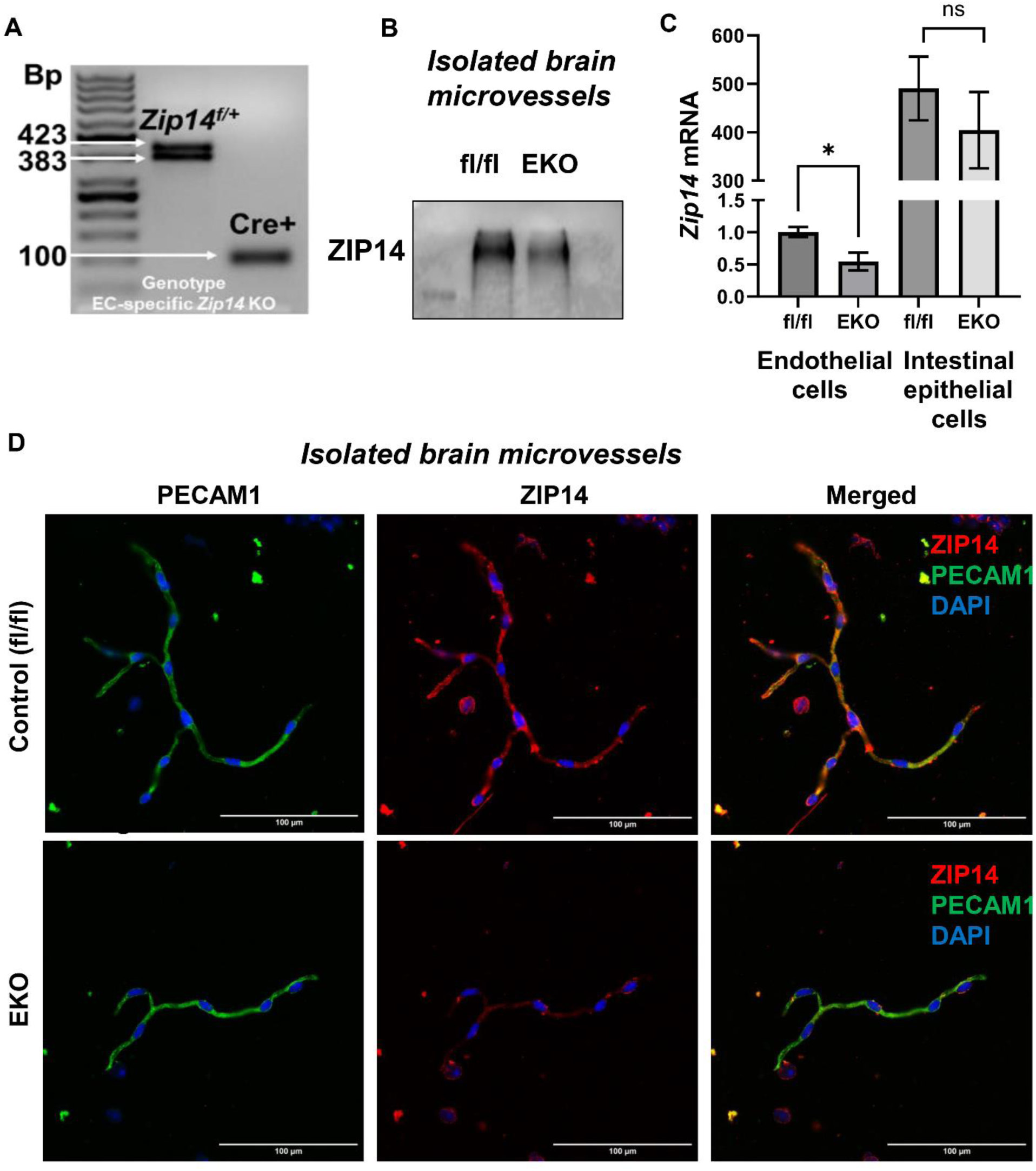
ZIP14 expression is reduced in EKO mice brain vessels. A) Representative PCR genotyping showing floxed (fl/fl) allele and Cre recombinase band in EKO mice. B) Western blot of ZIP14 protein expression in isolated brain microvessels from fl/fl control and EKO mice. C) *Zip14* mRNA expression in primary endothelial cells and intestinal epithelial cells isolated from fl/fl and EKO mice (n=3, mean±SEM, *p<0.05, Student’s t-test). D) Representative immunofluorescent images of ZIP14 (red) and PECAM1 (green) in isolated brain microvessels from fl/fl and EKO mice. Scale bars indicate image magnification.

To better understand how metal transporters interact within our ZIP14 knockout, we also explored the expression of other relevant metal transporters in isolated brain microvessels. Studies have shown that ZIP8 and ZnT10 are also present in endothelial cells, where they function in Mn uptake and export, respectively (39, 40, 46). In our study, we found that there is no compensatory upregulation of ZIP8 in EKO brain microvessels after Mn exposure (*Supplementary Figure 3)*.

However, ZnT10 expression was reduced, likely due to the lower intracellular Mn levels in EKO cells.

### EKO mice have greater brain Mn accumulation

Previous studies show whole-body *Zip14* knockout mice (WBKO) have brain Mn accumulation and subcutaneous injections of radioactive tracer ^54^Mn resulted in greater ^54^Mn levels in the brain of WBKO mice, suggesting ZIP14 is not required for brain Mn uptake (31). To see if EKO mice display similar phenotypes, we assessed brain Mn levels in EKO mice after dietary Mn supplementation (Figure 6A). Indeed, MRI analysis and direct measurement of brain Mn concentrations revealed significantly greater Mn accumulation in EKO mice compared to controls (*Supplementary Figure 4*, Figure 6B). Additionally, we evaluated Mn, Zn, and Fe levels in peripheral tissues after supplementation. EKO mice showed elevated blood Mn levels but no consistent changes in tissue Zn or Fe levels across tissues (*Supplementary Figure 5*).

**Figure 6.**
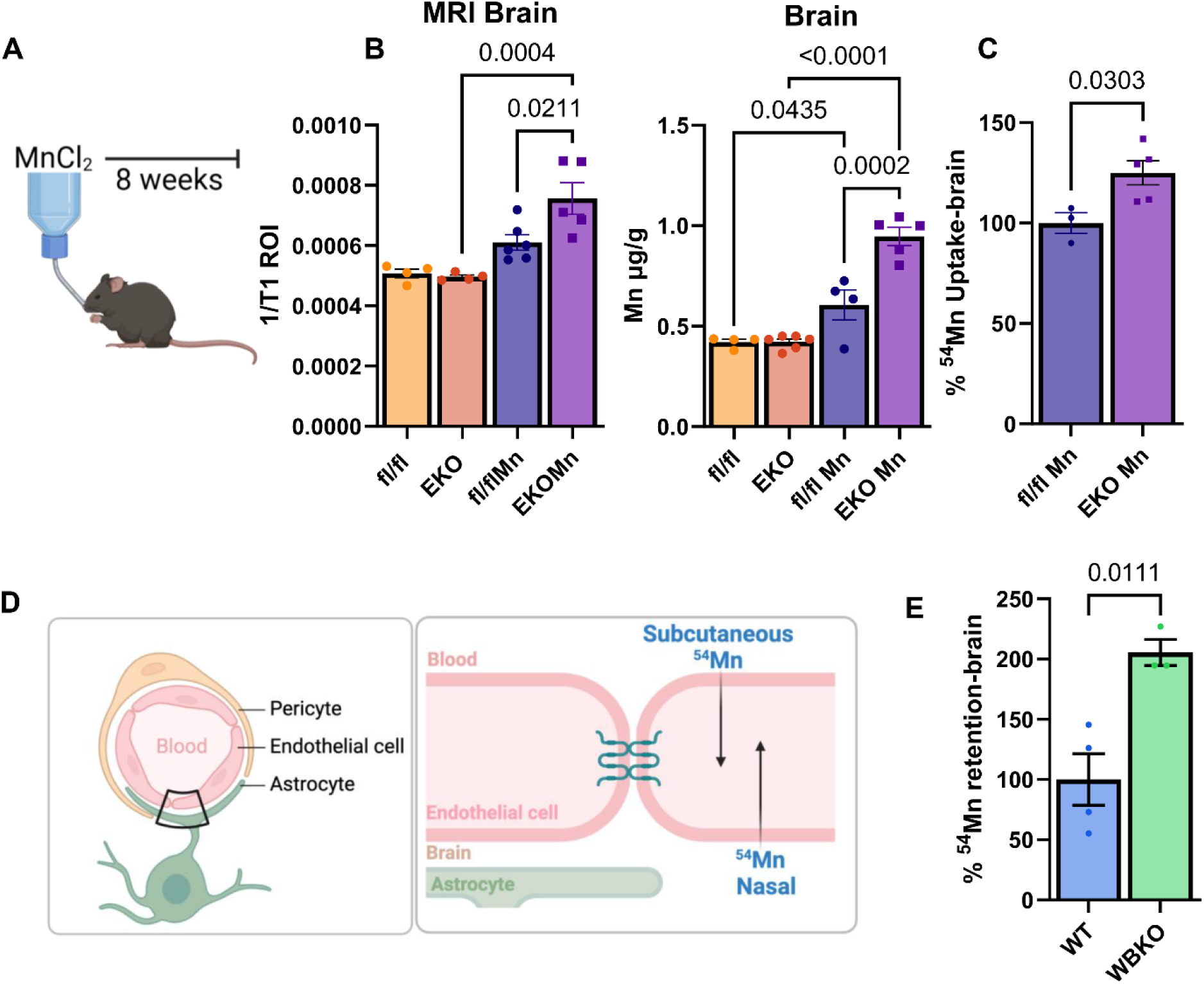
EKO mice accumulate more Mn in the brain and the dual delivery experimental system. A) Schematic showing a timeline of Mn supplementation in drinking water or vehicle control water for 8 weeks in fl/fl and EKO mice. B) Brain Mn accumulation as shown by T1-weighted MRI and total brain Mn levels measured by MPAES in digested brain tissue from fl/fl and EKO mice after supplementation (n=5). C) %^54^Mn uptake in the brains of fl/fl and EKO mice after subcutaneous injection of tracer ^54^Mn following Mn supplementation. D) Schematic of the dual delivery system, illustrating subcutaneous and nasal delivery routes for 54Mn tracer. E) %^54^Mn retained in the brain of WT (n=4) and whole-body ZIP14 knockout (WBKO; n=3) mice, measured 1h after nasal ^54^Mn delivery. Data represented as mean±SEM (Student’s t-test).

Similar to WBKO mice, subcutaneous ^54^Mn injection also resulted in increased brain ^54^Mn uptake in the EKO mice although to a lesser degree (Figure 6C) (31). Collectively, these findings suggest that ZIP14 is not essential for brain Mn uptake. No notable differences were observed in ^54^Mn uptake in other tissues (*Supplementary Figure 6*).

### ZIP14 facilitates Mn efflux from the brain

To investigate the direction of ZIP14-mediated Mn transport *in vivo*, we used a dual delivery system where we administered ^54^Mn via two routes, nasal delivery or subcutaneous injections (Figure 6D). Nasal delivery of Mn is well-established and in addition to being a good model of brain-to-blood direction, bypassing the BBB, it can be a good model for inhalation of airborne toxic metals (47–53).To validate the dual-delivery system, we first tested nasal delivery in WBKO mice and found that WBKO mice had two-fold greater ^54^Mn retained in the brain compared to wild-type mice (Figure 6E). From a previous study, we know that WBKO mice also have greater brain ^54^Mn after subcutaneous injections (31). These WBKO results suggest that ZIP14 could be required for Mn elimination but not critical for Mn uptake into the brain.

With our EKO model, we applied the dual-delivery system. In the steady state condition (i.e. without Mn supplementation), ^54^Mn retention in the brain was found to be higher in EKO mice compared to fl/fl mice (Figure 7A). This was true at 1h and 24h after the nasal ^54^Mn delivery; however, the effect is lost after 72h, suggesting that ZIP14 may play a more prominent role in Mn clearance during the early phase of elimination. The observed results indicate that the deletion of ZIP14 partially impairs Mn elimination from the brain, particularly during the first 24 hours. By contrast, subcutaneous injections showed no significant difference in brain ^54^Mn uptake between EKO and fl/fl control mice (Figure 7B). This indicates that ZIP14 is not directly involved in the movement of Mn from the blood to the brain, as deletion of ZIP14 did not change Mn transport in the blood-to-brain direction. Additionally, ^54^Mn uptake in peripheral tissues at 1, 24, and 72 hours post-nasal delivery showed no consistent differences between genotypes; however, there was increased ^54^Mn uptake in EKO peripheral tissues 3 hours post-subcutaneous injection (*Supplementary Figure 7-8)*. The increase in EKO peripheral tissue could be due to impaired initial ZIP14-mediated Mn elimination from the blood, leading to prolonged presence of ^54^Mn in the blood and subsequently increasing uptake into tissues via other transporters over time.

**Figure 7.**
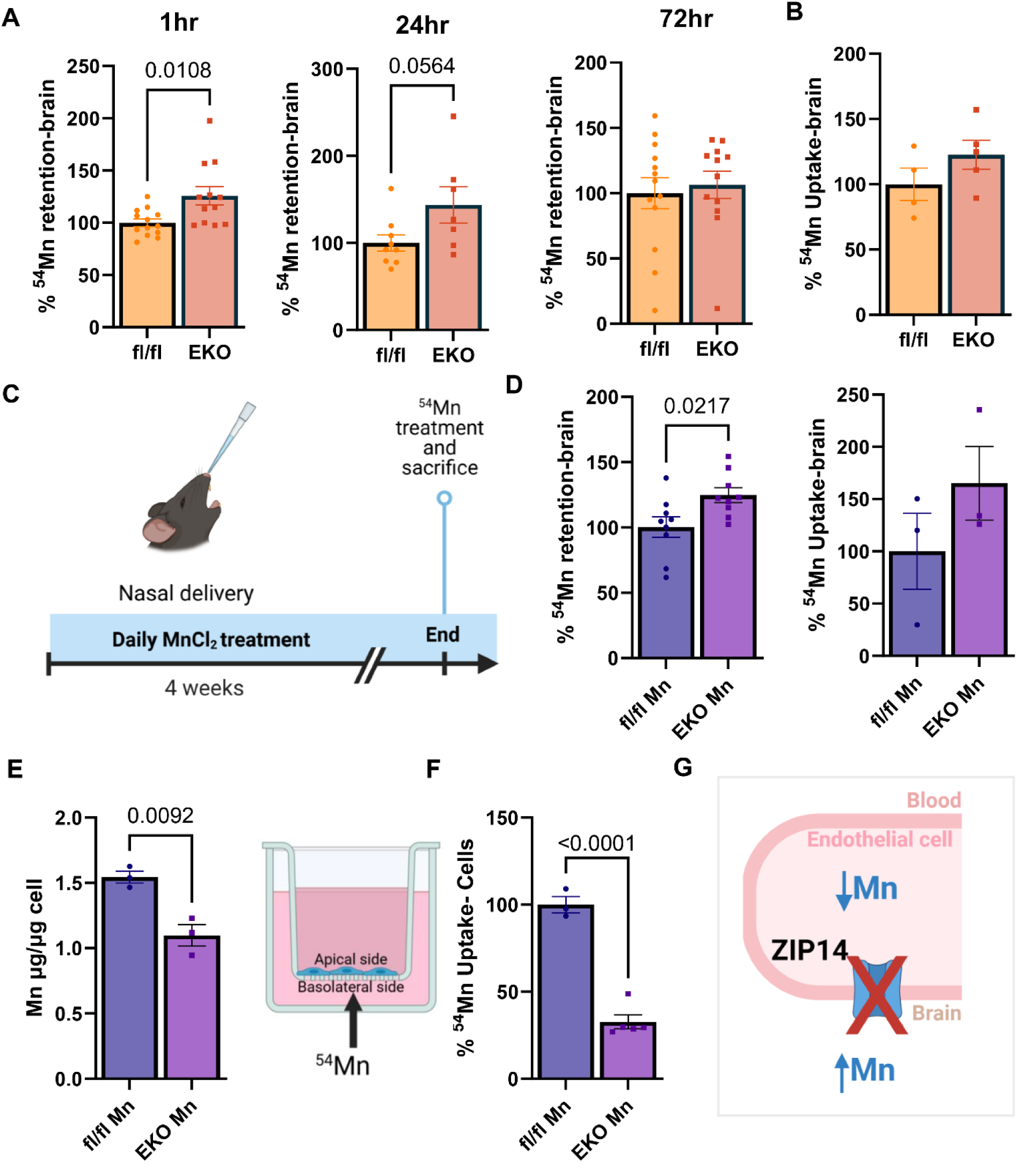
Greater brain Mn retention in EKO mice at steady state and after subchronic Mn supplementation. A) %^54^Mn retained in the brains of fl/fl and EKO mice at steady state (no Mn supplementation, measured at 1hr (n=13), 24hrs (fl/fl: n=7, EKO=9), and 72hrs (n=13) after nasal delivery. B) %^54^Mn uptake in the brain after subcutaneous injection of ^54^Mn in fl/fl and EKO mice at steady state (fl/fl: n=4, EKO=5). C) Diagram illustrating subchronic nasal Mn supplementation for 4 weeks, followed by tracer ^54^Mn delivery via nasal or subcutaneous routes (dual delivery system). D) %^54^Mn retained in the brain after nasal ^54^Mn delivery (n=9) and %^54^Mn uptake after subcutaneous injections in fl/fl and EKO mice (n=3) following subchronic nasal Mn supplementation. E) Mn levels in primary endothelial cells isolated from fl/fl and EKO cells following treatment with 300μM MnCl_2_ for 72hrs (n=3). F) Transwell ^54^Mn transport assay in primary fl/fl and EKO endothelial cells, seeded on Transwell inserts. ^54^Mn was added to the basolateral compartment and %^54^Mn uptake by cells was measured after 24hrs (fl/fl: n=3, EKO=5). G) Diagram depicting Mn accumulation in the brain and lowered Mn levels in the endothelial cell in the absence of ZIP14. Data are presented as mean±SEM, Student’s t-test.

Consistent with the steady state model, when EKO and fl/fl mice underwent subchronic nasal Mn supplementation for 4 weeks (Figure 7C), ^54^Mn was retained significantly more in EKO mice after nasal delivery, but again no significant difference was seen for subcutaneous injections (Figure 7D). Additionally, ^54^Mn uptake in peripheral tissues after dual delivery showed no significant differences between genotypes (*Supplementary Figure 9)*. Together, these findings support the hypothesis that ZIP14 plays a critical role in Mn elimination from the brain *in vivo*. Consequently, the absence of ZIP14 will lead to brain Mn accumulation.

To further validate the *in vivo* findings, we isolated and cultured primary endothelial cells from EKO and fl/fl mice. Following Mn treatment in cell media, we observed significantly lower Mn levels in EKO primary endothelial cells (Figure 7E). To assess directional Mn transport, fl/fl and EKO primary cells were seeded onto Transwell inserts, and once they reached appropriate TEER levels, ^54^Mn was added to the basolateral compartment. EKO cells had significantly lower levels of ^54^Mn (Figure 7F). Together, these results suggest that *Zip14* knockout impairs Mn uptake in the basolateral-to-apical direction, leading to decreased Mn levels within endothelial cells and contributing to Mn retention in the brain (Figure 7G). These findings support our *in vivo* data showing that ZIP14 is localized on the basolateral membrane and plays a key role in basolateral uptake of Mn, thereby facilitating Mn efflux from the brain.

#### ZIP14 overexpression upregulates Mn transport in the basolateral-to-apical direction

To further confirm the directionality of ZIP14-mediated Mn transport, we used hCMEC/D3 endothelial cell line with overexpression of ZIP14 (OE; Figure 8A). With increasing Mn treatment, OE cells have significantly greater Mn levels than WT cells (Figure 8B). Additionally, we observed that endogenous ZIP14 expression also increases with Mn exposure in wild-type (WT) hCMEC/D3 cells (*Supplementary 10*). Furthermore, when ^54^Mn was given to WT and OE cells, we found significantly greater ^54^Mn uptake by OE cells (Figure 8C). Similar to the primary endothelial KO cells, we seeded WT and OE cells on Transwell inserts and ^54^Mn was only added to the basolateral compartment. No difference was observed at 0.5h; however, by 24h, ^54^Mn levels were significantly lower in the basolateral compartment and correspondingly higher in the apical compartment (Figure 8D,F), with increased ^54^Mn accumulation in OE cells (Figure 8E) Together, these findings support the role of ZIP14 in mediating Mn transport in the basolateral-to-apical direction, highlighting its contribution to Mn efflux from the brain.

**Figure 8.**
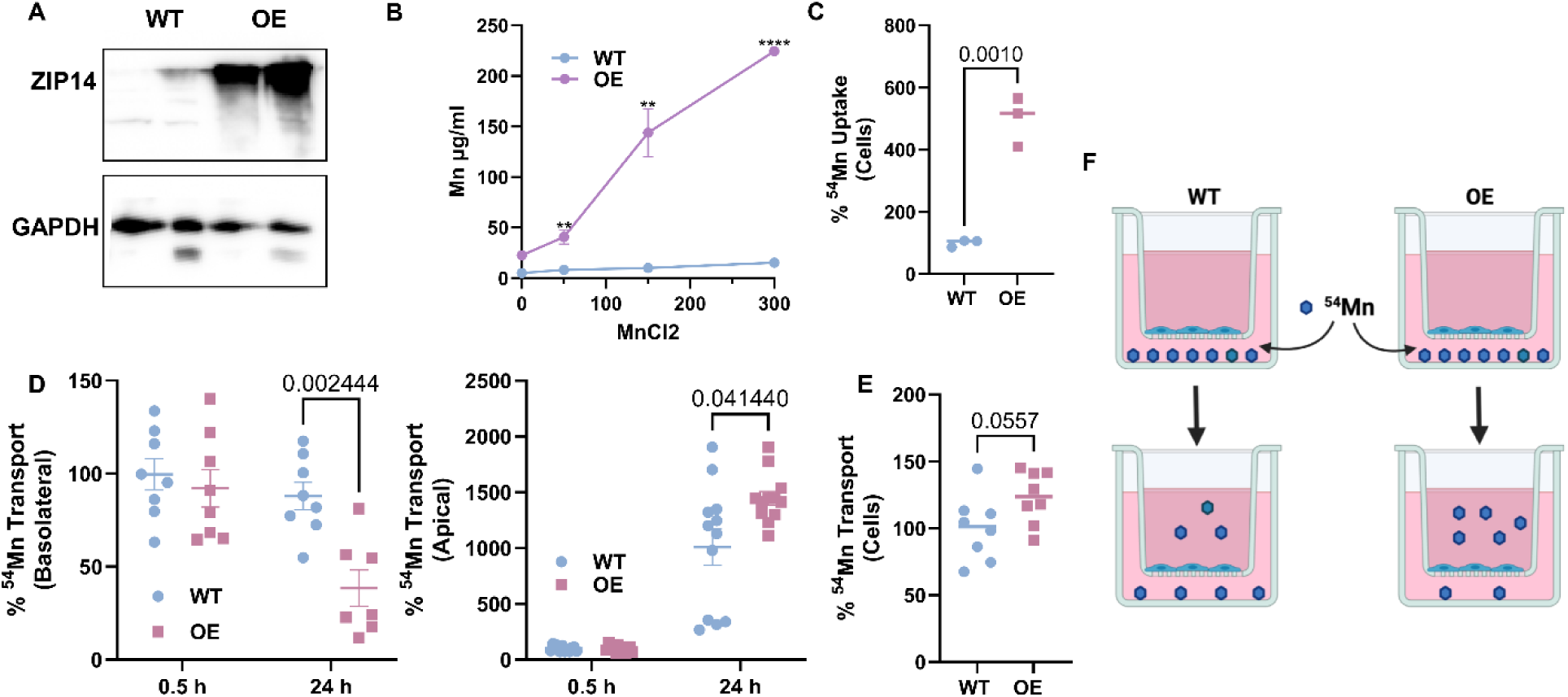
Overexpression of ZIP14 increases basolateral to apical transport of Mn. A) Western blot showing ZIP14 overexpression in hCMEC/D3 overexpressed (OE) cells compared to wild-type (WT) controls. B) Total Mn levels in WT and OE hCMEC/D3 cells after 0, 50, 150, 300μM MnCl_2_ treatment for 72hrs. (n=3, **p<0.01, ****p<0.0001). C) %^54^Mn uptake into WT and OE cells after ^54^Mn incubation for 24hrs (n=3). D-F) Transwell ^54^Mn transport assay: WT and OE cells are grown on Transwell inserts and ^54^Mn given to the basolateral side. Media samples were collected from apical and basolateral compartments at 0.5h and 24hrs, and cells were harvested at 24hrs. D) %^54^Mn measured in the basolateral and apical media at 0.5hrs and 24hrs of WT and OE cells (n=8). E) %^54^Mn retained in WT and OE cells after 24hrs (n=8). F) Schematic summarizing transport direction: After ^54^Mn addition to the basolateral compartment, OE cells had less ^54^Mn in the basolateral and more in the apical. Data are presented as mean±SEM (Student’s t-test).

## Discussion

Our study provides evidence that endothelial ZIP14 facilitates Mn efflux from the brain. We showed *in vivo* ZIP14 expression in brain endothelial cells and observed its upregulation in response to Mn. Using expansion microscopy, we noted that Mn exposure shifted ZIP14 localization from an equal apical-basolateral distribution to predominantly basolateral, supporting its role in facilitating brain Mn efflux. Using our endothelial-specific knockout mouse model with nasal ^54^Mn delivery and complementing *in vitro* ^54^Mn transport studies, we demonstrate that endothelial ZIP14 predominantly mediates basolateral-to-apical Mn transport, eliminating Mn from the brain, and its deletion results in brain Mn accumulation.

Our *in vivo* imaging showed ZIP14 expression in brain endothelial cells, supporting previous *in vitro* findings in the human brain microvascular endothelial cell line (hBMVEC) (54); however, its expression had not been validated *in vivo* or in other brain endothelial cell lines. We also demonstrated ZIP14 protein expression in isolated brain microvessels. A recent study by McCabe and Zhao (2024) showed *Zip14* gene expression in brain microvessels but did not detect any ZIP14 protein expression. The disparity in protein expression could be due to the expected ZIP14 molecular size. In our study, we detected ZIP14 protein at ∼250kb, whereas McCabe and Zhao (2024) focused only on the 50kb band size. Although ZIP14 has a predicted molecular mass of 58kb, Coffey and Knutson (2016) observed it at ∼130kb. Our previous work and other studies have consistently shown ZIP14 bands at ∼250kb, including in the brain endothelial cell line (54), and with a clear absence in *Zip14* knockout mice (31, 56, 57). Therefore, the higher molecular weight of ZIP14, likely due to oligomer formation, may have been overlooked in their study.

We also observed that its expression increased in response to Mn exposure, which partially aligns with the results reported by Steimle et al. (2019). The authors observed a 2-fold increase in ZIP14 expression in the plasma membrane without any change in the total protein content. The disparity may be due to the different Mn concentrations used. In their study, hBMVECs were incubated in 200 nM MnCl_2_ for 3.5h whereas we incubated hCMEC/D3 cells in 50-300 µM MnCl_2_ for 72 hours.

In zebrafish larvae, the overall *slc39a14* transcript level was seen to increase after 500µM MnCl_2_ exposure for 24 hours (17). In contrast, in hepatocytes, ZIP14 is degraded in response to Mn exposure (45). The authors showed that Mn induces downregulation of ZIP14 via the lysosomal degradation pathway and with that, reduced uptake of Mn. The authors suggested that this is a cryoprotective response to limit the uptake of excess Mn. This cryoprotective method does not seem to be present in brain endothelial cells.

Tight junctions between BBB endothelial cells restrict the passage of large molecules, requiring metal ions like Mn to rely on specialized transporters to enter or exit the brain (58). BBB endothelial cells differ from peripheral endothelial cells due to their polarized structure, which results in different protein distributions between its apical versus basolateral side (59). Given the polarized and extremely thin nature of BBB endothelial cells, determining the localization of metal transporters on the apical or basolateral membranes has remained challenging in the field. To overcome this limitation, we used expansion microscopy, a technique that allowed us to physically expand brain sections 3-fold, significantly improving the resolution of endothelial cells surrounding blood vessels. Combined with light sheet fluorescence microscopy, this approach allowed direct *in vivo* visualization of transporter localization (60–62). Using this technique, we reported a shift of ZIP14 distribution from the apical to basolateral surface of BBB endothelial cells following Mn supplementation. Our results align with those reported by Steimle et al. (2019), who found ZIP14 expression in both membranes *in vitro*, with significantly greater expression on the basolateral side. However, their observations were made without Mn supplementation *in vitro*, whereas ours were observed following Mn exposure *in vivo*. This discrepancy may reflect inherent differences between *in vitro* and *in vivo* systems, emphasizing the value of direct visualization methods with expansion microscopy.

Our subcutaneous ^54^Mn injections showed a slight, though non-significant, increase in brain ^54^Mn uptake in EKO mice, with and without nasal supplementation—a trend also seen in dietary Mn-supplemented EKO mice. This increase may not reflect increased uptake but rather reduced Mn elimination from the brain after *Zip14* deletion. To further validate this, we demonstrated that *in vitro* ZIP14 knockout reduced cellular ^54^Mn uptake from the basolateral side, while ZIP14 overexpression increased basolateral-to-apical transport and cellular ^54^Mn uptake from the basolateral side. These results are consistent with the observations by Steimle and group (2019), who also reported that ZIP14 knockdown significantly reduced ^54^Mn uptake from the basolateral side. Moreover, they reported that wild-type cells exhibited greater ^54^Mn uptake at the basolateral side than at the apical side, suggesting that BBB endothelial cells may preferentially transport Mn from the brain-facing side to facilitate brain Mn elimination, consistent with our findings.

One limitation of our study is that we did not include behavioral assessments of our EKO mouse model. Individuals with *SLC39A14* mutations (17–26) and WBKO mice (27–32) exhibit motor and/or cognitive deficits, suggesting a link between ZIP14 function, Mn accumulation, and neurological outcomes. Although behavioral analysis is beyond the scope of this study, future work should include comprehensive spatial brain Mn accumulation and behavioral tests for motor and cognitive functions in this animal model.

In conclusion, our study demonstrates that BBB endothelial ZIP14 plays an important role in brain Mn clearance. With Mn exposure, ZIP14 shifts to greater localization on the basolateral membrane and primarily facilitates brain Mn efflux *in vivo*. These findings provide mechanistic insight into how the brain protects itself against Mn overload and why ZIP14 mutations in humans result in brain Mn accumulation and symptoms of Parkinsonism.

## Materials and Methods

### Animals

Mice were fed *ad libitum* a commercial-chow type diet (PicoLab® Rodent Diet 20 containing 82mg/kg Mn) and water containing no detectable Mn. All mice were housed in a specific pathogen-free facility and received the same standard husbandry. Protocols were approved by the Cornell University Institutional Animal Care and Use Committees.

Transposagen Biopharmaceuticals generated floxed *Zip14* mice on the C57BL/6 background. Founder *Zip14*^flox/+^ (LoxP in introns 4 and 8) mice were bred to obtain *Zip14*^flox/flox^ mice (fl/fl). fl/fl mice were crossed with (B6.Cg-Tg[Tek-cre]1Ywa/J-Jackson Lab stock # 008863) mice to create *Tie2*-expressing cell-specific *Zip14* KO mice (EKO). Genotyping of EKO mice was performed from DNA extracted from mice ear punches, using the forward primer 5’-GCG GTC TGG CAG TAA AAA CTA TC-3’ and the reverse primer 5’-GTG AAA CAG CAT TGC TGT CAC TT-3’, amplifying a 100bp product. fl/fl mice were used as controls. Mice from both sexes were used in this study.

### Cells

Primary brain endothelial cells from adult fl/fl and EKO mice were isolated with protocols modified from previous studies (63, 64). Briefly, mice brains were collected, meninges, olfactory bulbs and cerebellums were removed. Brains were cut into pieces and homogenized using a Teflon Dounce homogenizer. Myelin was separated and removed by suspension of the pellet in 18% dextran and centrifugation. The resulting pellet contained vessels and was digested with collagenase/dispase. The product was centrifuged and seeded onto collagen type I rat tail-coated plates. The cells were cultured in DMEM-F12 with 20% fetal bovine serum, 10% astrocyte-conditioned media, 20 USP/ml heparin sodium (Sagent), 2mM GlutaMAX™, 1% penicillin-streptomycin and 0.2% Normocin® (InvivoGen). The first two days, they were given 4µg/ml puromycin (64). The primary cells were passaged once before any assays were performed.

To obtain pure primary endothelial cells, we modified our existing protocol to obtain a single-cell suspension, followed by a positive selection of endothelial cells (65–67). Briefly, we added an extra 50-minute digestion step using collagenase II before the myelin separation. After the collagenase/dispase digestion step, we used CD31 microbeads (Miltenyi Biotec 130-097-418) and MS MACS Columns to select mouse brain endothelial cells. Pure cells were seeded on collagen-coated 24-well plates.

We also used the immortalized human brain microvascular endothelial cell line hCMEC/D3 obtained from Dr. Claudia Fischbach-Tesch with Dr. Babette B. Weksler’s permission. Stably ZIP14 overexpressing hCMEC/D3 cells were custom-made via transduction with the *ZIP14/SLC39A14* overexpression lentivirus by Applied Biological Materials Inc. (abm). Cells were seeded on collagen type I rat tail (Millipore Sigma) coated cell culture dishes and cultured in EBM-2 media (Lonza) supplemented with EGM-2 SingleQuots (Lonza), 10% astrocyte conditioning media (ACM, ScienCell) and 1% penicillin-streptomycin (ThermoFisher). All cells were maintained at 37 °C in a humidified atmosphere of 95% air and 5% CO2.

### Mn exposure models

*In vivo.* To model subchronic nasal Mn exposure, mice were given 4μl nasal delivery of 30mg/kg MnCl_2_ daily for 4 weeks as previously described (68). Briefly, mice were anesthetized via isoflurane inhalation, and droplets of MnCl_2_ were administered to the left or right nostril, alternating daily.

To model chronic dietary Mn exposure, fl/fl and EKO mice were given 7.2g MnCl_2_•H_2_O in their drinking water for 8 weeks (Mn consumption is ∼20 mg/kg body weight/day). Several previous studies demonstrated Mn neurotoxicity at similar concentrations (69–75).

*In vitro.* To assess ZIP14 expression and Mn concentrations following Mn exposure, primary endothelial cells and hCMEC/D3 cells were treated with various concentrations of MnCl_2_ in media for 72h.

### ^54^Mn transport

*In vivo*. Animals were anesthetized via isoflurane inhalation. 4µl (0.2-0.35µCi) of ^54^MnCl_2_ (Eckert & Ziegler) was administered into the left or right nostril of the mouse. 1h, 24 h, or 72h later, mice were sacrificed, then tissue samples were collected and weighed for normalization. We observed the highest brain ^54^Mn levels at 1h, consistent with a previous study (76). Based on this, we selected the 1h time point for our ^54^Mn nasal delivery experiment after nasal Mn supplementation.

For nasal delivery, brain tissue radioactivity counts per minute (cpm) were calculated using the following equation 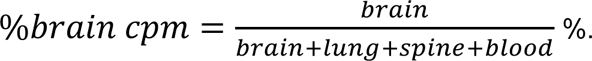

This equation accounts for the potential movement of ^54^Mn into the nasal blood vessels, cerebrospinal fluid (indicated by the spine cpm), or leakage into the lung (*Supplementary Figure 11*). The resulting percentage represents the portion of radioactivity remaining in the brain after 1, 24, or 72h, compared to the total amount absorbed. Note that the total amount absorbed does not equal the total amount administered, as a variable volume of liquid could remain in the nasal canal.

For subcutaneous injections, mice were fasted for 4h and then injected subcutaneously with 100µl of ^54^MnCl_2_ (20 µCi/ml). Mice were sacrificed 3h post-injection, then tissue samples were collected and weighed. For both nasal and subcutaneous delivery, radioactivity was measured with the gamma counter (PerkinElmer Wallac Wizard 3" 1480) and normalized to tissue weight.

*In vitro*. Primary endothelial cells and hCMEC/D3 cells were seeded on 0.4μm pore size and 3.0μm Transwell membranes, respectively. Transendothelial electrical resistance (TEER) measurements were recorded to confirm monolayer confluency, ensuring that cells reached appropriate resistance values before experimental use, primary brain endothelial cells reaching 60Ω•cm^2^ (77) and hCMEC/D3 cells reaching 35Ω•cm^2^ (78). Cells were pretreated with 300μM MnCl_2_ for 72h. Following pretreatment, 0.18μCi/ml ^54^Mn was added to serum-free endothelial media with 50μM MnCl_2_ and given to the basolateral compartment, whereas serum-free endothelial media was added to the apical compartment. Media from both sides were collected at 0.5h for radioactivity measurement and then replaced with fresh radioactive media for the next 24hs. At endpoint, media was removed, and cells were washed with wash buffer (0.9% NaOH, 10mM EDTA, 10mM HEPES), then solubilized with 0.2% SDS and 0.2M NaOH for 1h. Radioactivity, counts per minute (cpm), was measured for both cell lysates and media using a gamma counter. Protein concentration was quantified using BCA assay, and cpm values were normalized to total protein.

### Purification of mouse brain microvessels

Animals were anesthetized via isoflurane inhalation and underwent intracardiac perfusion with 20 ml of ice-cold PBS to remove blood from the brain vessels. After removal of the choroid plexus, olfactory bulbs, and cerebellum, the brain was homogenized. First, the brain was cut into smaller (∼2mm) pieces using scissors and then homogenized with an automatized Dounce homogenizer (63). The homogenate then underwent centrifugation with 18% (w/v) dextran before being resuspended and passed through a 20 µm-mesh filter. The purified brain vessels were present on the filter and plated on poly-L-lysine-coated coverslips for immunofluorescent staining.

### Endothelial tubule formation

Endothelial tubule formation was carried out by seeding primary fl/fl endothelial cells on the Matrigel (50µl/well) of a 96-well plate and incubated at 37°C for 7h. Six phase contrast images were taken for each well using the BioTek Lionheart FX Automated Microscope.

### Immunofluorescence

Animals were anesthetized via isoflurane inhalation and underwent intracardiac perfusion fixation with 20ml of ice-cold PBS followed by 20ml ice-cold 4% paraformaldehyde (PFA). Brains were removed, fixed in 4% PFA overnight, and then transferred to 30% sucrose until they sank to the bottom of the solution. The brains were embedded in OCT (4583, Sakura), frozen, and cryosectioned at a thickness of 15μm. Primary mouse endothelial cells, hCMEC/D3 cells, and Matrigel tubules were fixed with 2% PFA at room temperature for 20min.

Both brain sections and cells were permeabilized with 0.3% Triton X for 10min and then blocked with 10% normal goat serum, 3% bovine serum albumin (BSA), and 0.1% Triton X in PBS for 1 h at room temperature. Following the blocking step, samples were incubated at 4℃ overnight with the following primary antibodies: rabbit anti-mouse ZIP14 antibody 10µg/ml (custom-made by Genscript Antibodies) (57, 79), PECAM1 1:100, and rabbit anti-human ZIP14 antibody 1:200 (LSBio, LS-B7008). For immunofluorescent tubules, primary antibody incubation was performed at room temperature for 1h instead.

Secondary incubation was performed at room temperature for 1h with AlexaFluor594 goat anti-rabbit 1:1000 (Thermo Fisher Scientific A-11012), AlexaFluor488 goat anti-rat 1:500 (Thermo Fisher Scientific A-11005) secondary antibodies. Matrigel tubules were further incubated with DAPI solution, and the slides were mounted in Antifade Mounting Medium with DAPI (Vectashield). All images were taken with BioTek Lionheart FX Automated Microscope.

### Expansion microscopy

Following intracardiac perfusion with 4% PFA and overnight 4% PFA fixation, brains were embedded in 4% agarose and sectioned coronally at 100µm thickness with a vibrating blade microtome (Leica VT1200 S). Immunofluorescent staining was performed on free-floating brain sections (100µm) in 24-wells and a region containing blood vessels was selected and excised. Tissue was anchored with AcX (0.2 mg/ml) overnight at 37°C, embedded in the polyacrylate hydrogel and digested overnight with proteinase K (80). The sample was then expanded in 0.1X PBS to achieve 3-fold linear expansion and imaged on a lightsheet fluorescence microscopy (Zeiss Z.7). A 20 × water-immersion objective (20x/1.0 W Plan-Apochromat Corr DIC M27 75mm, RI = 1.33) was used for imaging.

Quantification was performed in ImageJ. Regions of interest (ROIs) were manually drawn around the apical and basolateral surfaces after identifying individual endothelial cells (*Supplementary Figure 1-2A)*. Mean ZIP14 fluorescence intensity was measured within each ROI. For overall ZIP14 intensity, values were normalized to fl/fl controls (set to 100%). For apical and basolateral measurements, all values were normalized to fl/fl apical intensity (set to 100%).

### Magnetic Resonance Imaging (MRI)

MRI images were acquired at the Cornell Magnetic Resonance Imaging Facility on a General Electric 3.0 Tesla MR750 scanner (Waukesha, WI) with an eight-channel flexible receive surface coil. Mice were induced with 4% isoflurane in oxygen for 5 min and then kept anesthetized with 2% isoflurane during MRI. Vital signs were monitored during MRI with a MouseOx Plus system by Starr Life Sciences Corp. (Oakmont, PA). The overall scan time was about 50 min per animal.

Approximate T1 maps were obtained from 14 transverse slices of the brain using a multiple repetition time spin echo sequence (41) (TR = 550 ms, 1000 ms, 2000 ms) with echo time = 12 ms, flip angle = 90 degrees, and parallel imaging acceleration (ARC) factor = 2, with the following geometric parameters: Field-of-view = 6 cm × 6 cm, acquisition matrix size = 192 × 192 (in-plane image resolution after zero filling to a 512 × 512 matrix = 0.1172 mm × 0.1172 mm), slice thickness = 1.2 mm. Each image was acquired with an NEX of 16, 8, and 4, depending on the TR. The frequency direction was A/P, and the bandwidth was 20.83 KHz. A customized Matlab script to perform voxel-wise nonlinear least squares fitting to the equation S_TR_=S_0_(1-e^-TR/T1^) was used to generate approximate parametric T1 maps for further statistical analysis.

ImageJ was used to manually trace ROIs encompassing the whole brain in each transverse MRI slice. Mean signal intensity was calculated for each slice and then averaged across slices to obtain a single mean intensity value per animal.

### Metal content measurements (metallomics)

Animals were sacrificed, and their tissues were collected, weighed, and digested in HNO_3_ overnight at 85°C. The small intestine was divided into proximal (10cm) and distal segments (remaining length). Digested samples were then diluted 1:1 in Milli-Q water and Mn, Fe, Zn concentration was measured using Microwave Plasma-Atomic Emission Spectrometry (MP-AES) at 403.076nm, 371.993 nm, 213.857 nm, respectively. The resulting values were normalized to tissue weight. Cells were harvested from 6-well or 24-well plates and then underwent digestion and dilution following the procedure described above. MPAES values for cells were normalized to protein levels obtained from total protein concentrations.

### RNA isolation and qPCR

Intestinal epithelial cells were isolated from animals by isolating intestinal segments and flushing the lumen. Intestinal segments were inverted over bamboo sticks and immersed in 15ml tubes with 1xPBS with 1.5 mM EDTA. The sticks were then twisted on ice every few minutes to release intestinal epithelial cells. After 20 mins, tubes were centrifuged for 5 min at 500xg. Samples were washed three times with 10ml cold PBS and centrifuged again to collect intestinal epithelial cells (57). Total RNA was isolated from 100µl of the pelleted cells.

Total RNA was isolated from pure brain endothelial cells grown in cell culture and isolated brain microvessels using Trizol. cDNA synthesis was performed with M-MLV reverse transcriptase and gene expression was measured with PowerTrack SYBR Green Master Mix using Applied Biosystems Quantstudio3. Primers we used in this study include: *Zip14* forward 5’-TTTCCCAGCCCAAGGAAG-3’, *Zip14* reverse 5’-CAAAGAGGTCTCCAGAGCTAAA-3’, *Zip8* forward 5’-GCCAAGCTCATGTACCTGTCT-3’, *Zip8* reverse 5’-AAGATGCCCCAATCGCCAA-3’, *Znt10* forward 5’-GGCCGTTACTCAGGCAAGAC-3’, *Znt10* reverse 5’-GCATGTTGAACGAGTCCGAGA-3’, *Gapdh* forward 5’-AGGTCGGTGTGAACGGATTTG-3’, *Gapdh* reverse 5’-GGGGTCGTTGATGGCAACA-3’. Data was normalized to *Gapdh*.

### Western Blot analysis

hCMEC/D3 WT and OE cells were seeded on 10cm dishes and harvested using non-denaturing lysis buffer (20mM Tris HCl pH 8.0, 137mM NaCl, 10% glycerol, 1% Triton X-100, 2mM EDTA), supplemented with protease and phosphatase inhibitors, as well as PMSF. Isolated brain microvessels were homogenized in a non-denaturing lysis buffer. BCA was performed to determine protein concentration and equal amounts of protein were loaded onto 10% SDS-PAGE gels and visualized using chemiluminescence (SuperSignal, Thermo Fisher) and digital imaging (Protein Simple). The following primary antibodies were used: mouse ZIP14 antibody 4µg/ml (custom-made by Genscript (81–84)), human ZIP14 1:1000 (LSBio, LS-C668142), and GAPDH 1:1000 (Cell Signaling, 14C10).

### Statistical analysis

All statistical analyses were performed in GraphPad Prism (version 9.5.1). Two-group comparisons will be performed with Student *t*-tests. Three or more group comparisons were achieved through 1-way ANOVA. Statistical significance is at P < 0.05. Data sets are presented as means ± SE.

## Acknowledgments

The research is funded by Cornell University Division of Nutritional Sciences, Funding to TBA, the Cornell University Center for Vertebrate Genomics (CVG) grant to TBA, and the Cornell Magnetic Resonance Imaging Facility (CMRIF) pilot grant to TBA. We thank Drs. Barbara Strupp, Nozomi Nishimura, and Martha Field for the critical discussions and data interpretation. We also thank Drs. Sumit Narayan Niogi and Henning Voss at the Cornell MRI facility for their contribution to developing the method for MRI.

## Data Availability

The authors confirm that the data supporting the findings of this study are available within the article [and/or] its supplementary materials.

## Author Contributions

TBA conceived and designed research; TBA, TLT, JZ, ZW performed experiments; TBA, YW, TLT, and JZ, analyzed data; TBA, YW, JZ, and TLT interpreted results of experiments; JZ prepared figures; TBA and JZ drafted the manuscript; TBA, JZ, TLT, YW edited and revised manuscript; TBA, JZ, TLT, YW and ZW approved the final version of the manuscript.

