## Supplemental Figures for "The Ins and Outs of Manganese: ZIP14 facilitates the efflux of excess manganese from the brain"

**Supplementary Figures**

Supplementary Figure 1 Quantification images for (no Mn treated) expanded brain with scale bar

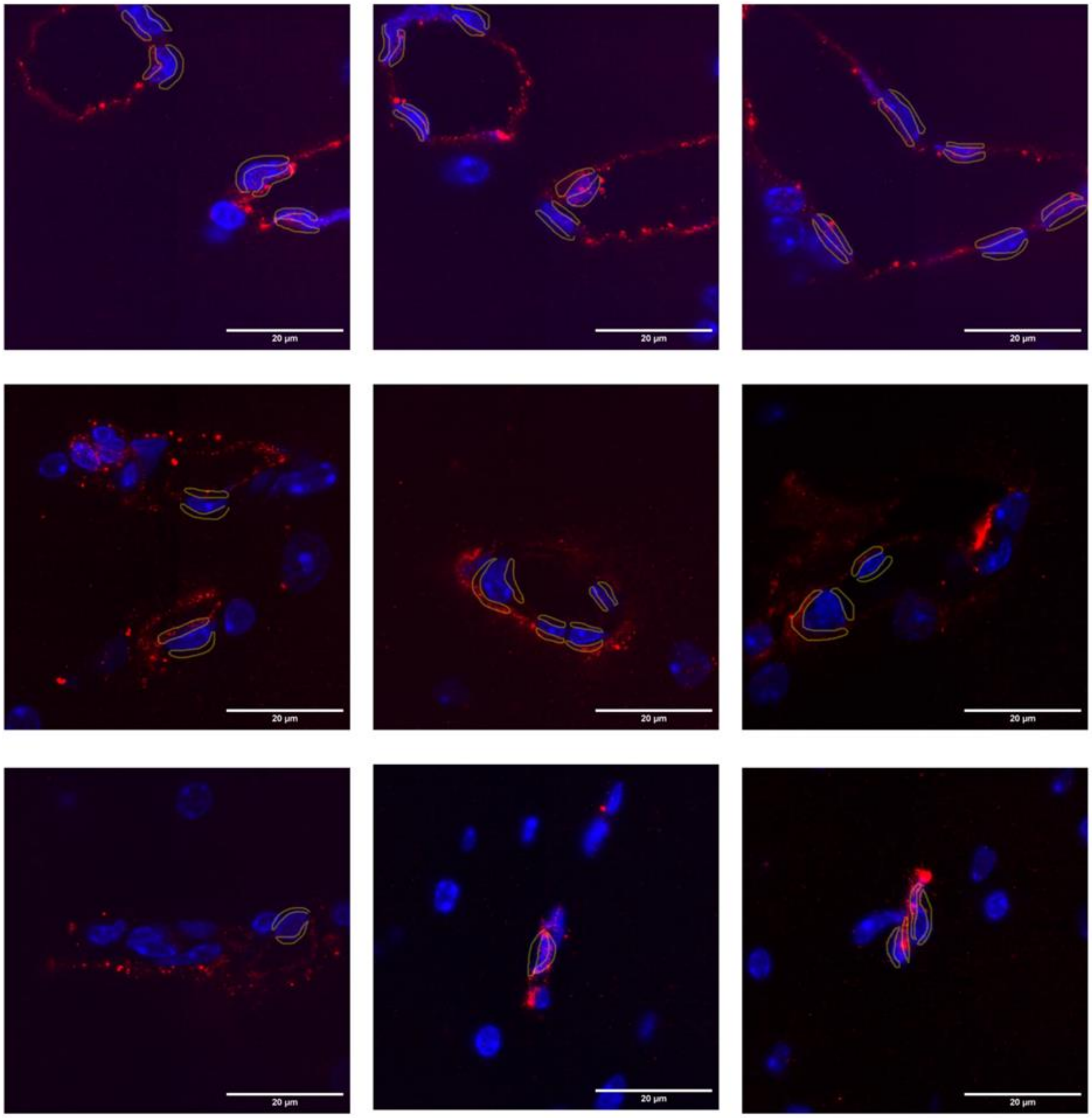


**Supplementary Figure 1.** Fluorescent images of blood vessels from control mouse brains without Mn supplementation. Quantification of ZIP14 intensity was performed in apical and basolateral areas of individual endothelial cells, as denoted from the yellow outlines. Scale bars indicate image magnification.

Supplementary Figure 2 Quantification images for Mn treated expanded brain with scale bar

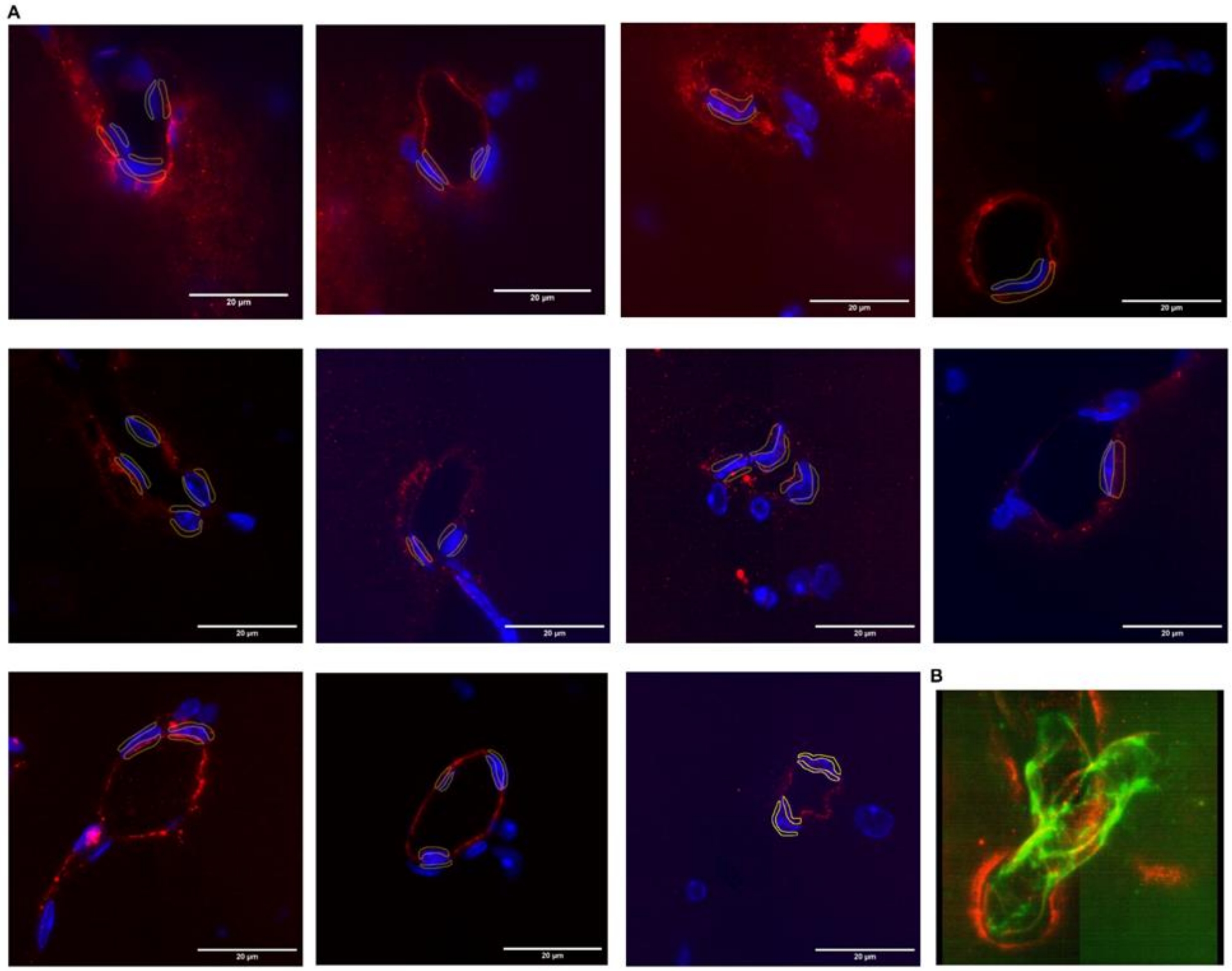


**Supplementary Figure 2.** Fluorescent images of blood vessels from control mouse brains after Mn supplementation. Quantification of ZIP14 intensity was performed in apical and basolateral areas of individual endothelial cells, as denoted from the yellow outlines. Scale bars indicate image magnification. B) Representative video of a 3D reconstruction from z-stacks of a brain vessel from fl/fl mice following Mn supplementation, highlighting ZIP14 (red) and PECAM1 (green) localization.

Supplementary Figure 3 *Zip8* and *Znt10* expression in blood vessels isolated from EKO brain.

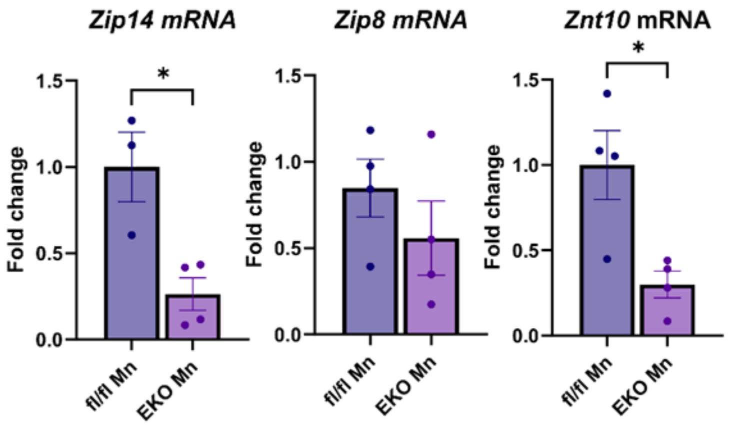


**Supplementary Figure 3**. *Zip14, Zip8,* and *Znt10* expression in EKO vessels. Brain microvessels were isolated from fl/fl and EKO mice after 4 weeks of nasal Mn supplementation. RNA was extracted and analyzed by qPCR for *Zip14,*

*Zip8,* and *Znt10* expression (n = 4). mRNA levels were normalized to *Gapdh*. Data are presented as mean±SEM, Student's t-test

Supplementary Figure 4 Tissue Mn, Zn, Fe levels after dietary Mn supplementation

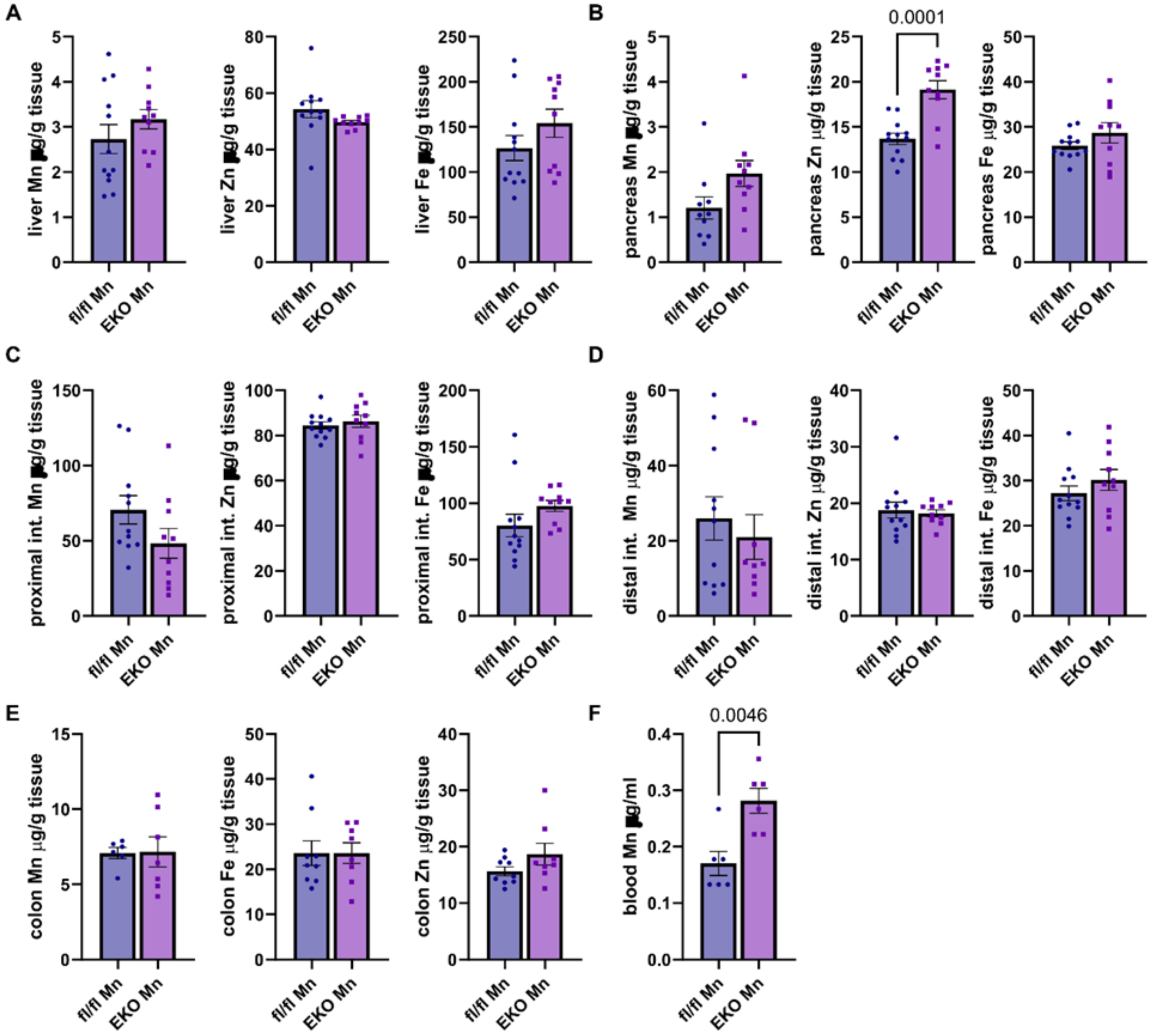


**Supplementary Figure 4.** Brain MRI images of fl/fl and EKO brains with and without Mn supplementation. Representative T1-weighted MRI images of brains from fl/fl and EKO mice after 8 weeks of dietary Mn

supplementation or vehicle control. MRI on anesthetized mice was performed on a General Electric 3.0 Tesla scanner (Waukesha, WI). Both data sets were analyzed with customized MATLAB software to process approximate T1 maps.

Supplementary Figure 5. Tissue %54Mn levels after subcutaneous injection following dietary Mn supplementation

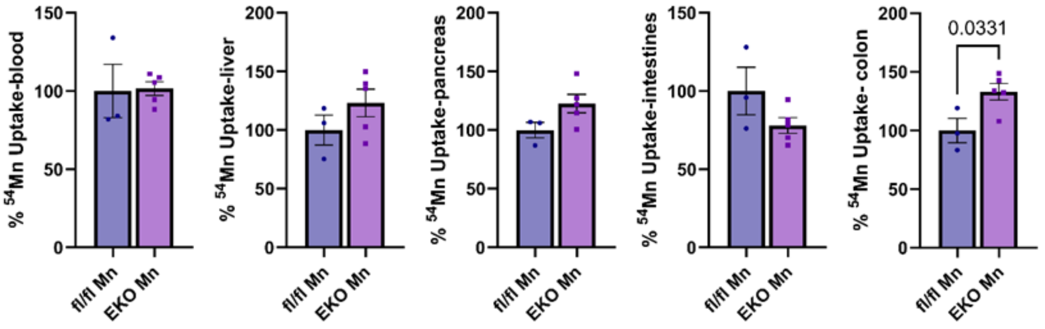


**Supplementary Figure 5.** Tissue Mn, Zn, and Fe levels after dietary Mn supplementation. fl/fl and EKO mice were given dietary Mn-supplemented drinking water for 8 weeks. At endpoint, A) liver, B) pancreas, C) proximal intestines, D) distal intestines, E) colon and F) blood were collected, digested with nitric acid, and Mn, Fe, Zn concentrations were measured using microwave plasma atomic emission spectroscopy (MPAES). Metal levels were normalized to tissue weight or blood volume. Note, only Mn levels were measured for blood due to the lack of blood volume. Data are presented as mean±SEM, Student's t-test.

Supplementary Figure 6. Brain MRI images of fl/fl and EKO brains with and without Mn suppelmentation

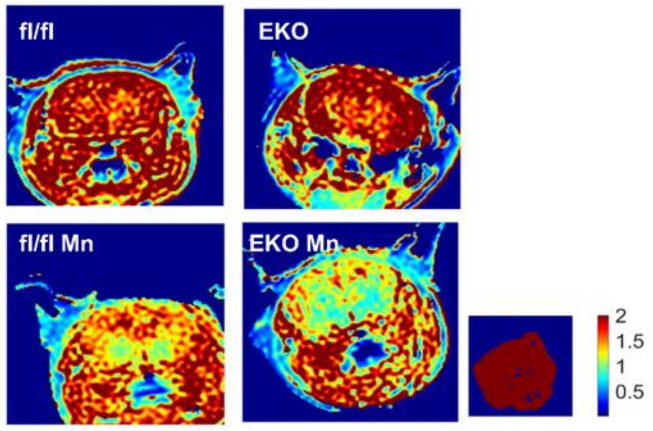


**Supplementary Figure 6.** Tissue %$^{54}$Mn uptake after subcutaneous injection following dietary Mn supplementation. fl/fl and EKO mice were injected with $^{54}$Mn subcutaneously after a 4h fast, following 8 weeks of dietary Mn

supplementation in water. 3h post-injection, tissues were collected, and radioactivity was measured with a gamma counter and normalized to tissue weight. EKO counts per minute (cpm) are presented as a percentage relative to fl/fl cpm. Sample sizes: Data are presented as mean±SEM, Student's t-test

Supplementary Figure 7. Tissue %54Mn uptake levels at steady state (no Mn supplementation) post-nasal delivery

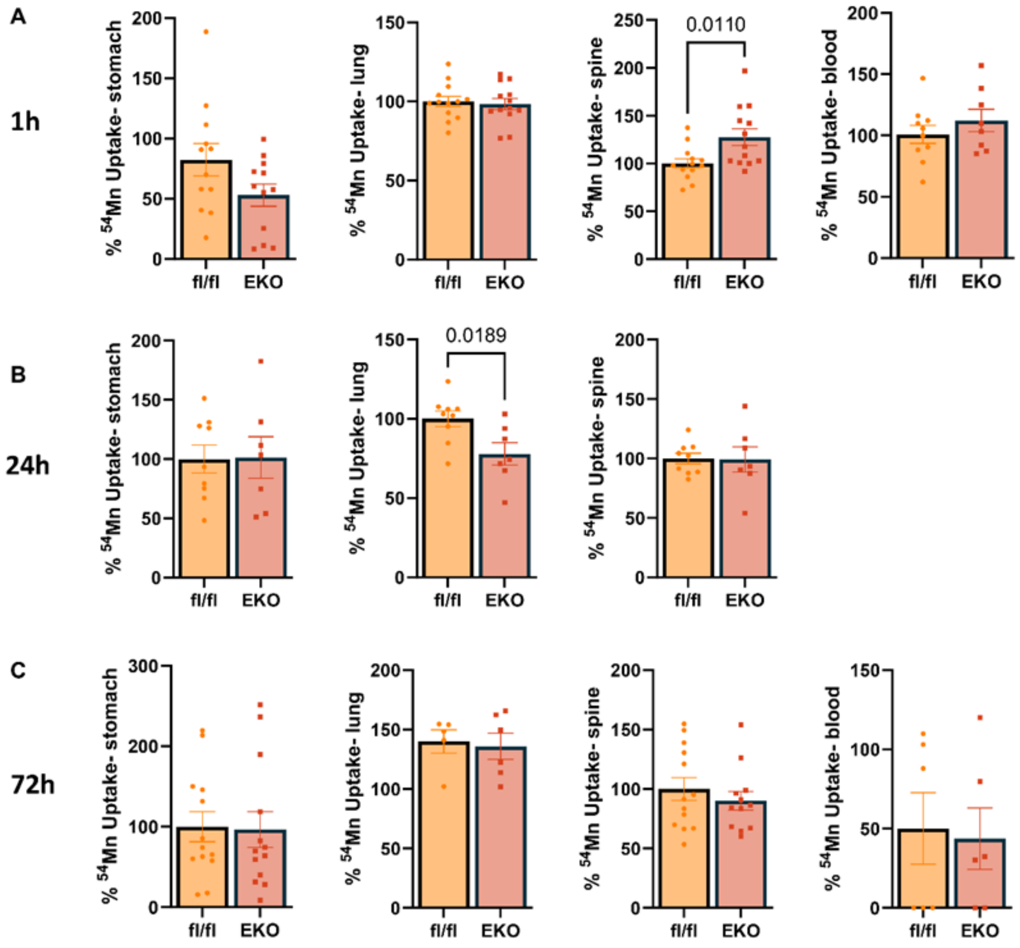


**Supplementary Figure 7.** Tissue %$^{54}$Mn uptake levels at steady state post-nasal delivery. l/fl and EKO mice received nasal $^{54}$Mn delivery under steady-state conditions (no Mn supplementation). Tissues were collected at three time points: A) 1h, B) 24h C) 72h. Tissue radioactivity (counts per minute, cpm) was measured using a gamma counter and normalized to tissue weight. EKO cpm values are presented as a percentage relative to fl/fl cpm. Data are presented as mean ± SEM; statistical significance was assessed using Student's t-test.

Supplementary Figure 8. Tissue %54Mn uptake levels post-subcutaneous injection at steady state (no Mn supplementation)

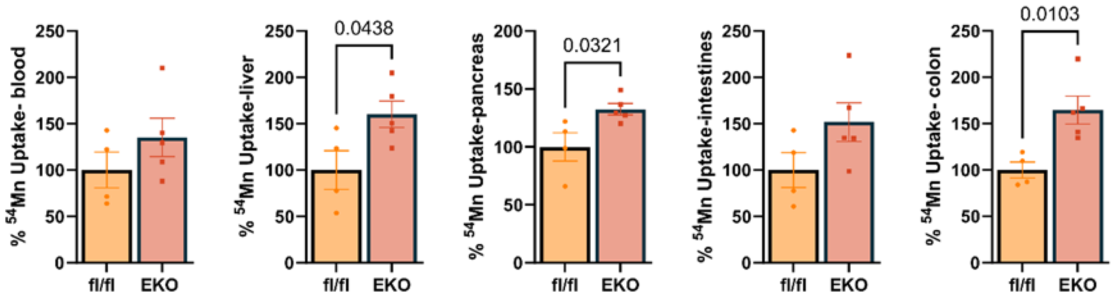


**Supplementary Figure 8.** Tissue %$^{54}$Mn uptake levels at steady state post-subcutaneous injections. fl/fl and EKO mice were injected subcutaneously with $^{54}$Mn under steady-state conditions (no Mn supplementation). Three hours post-injection, tissues were collected, and radioactivity (counts per minute, cpm) was measured using a gamma counter and normalized to tissue weight. EKO cpm values are presented as a percentage relative to fl/fl cpm. Data are presented as mean ± SEM; statistical comparisons were made using Student's t-test.

### Supplementary Figure 9. Tissue %54Mn uptake levels after dual delivery following Mn supplementation

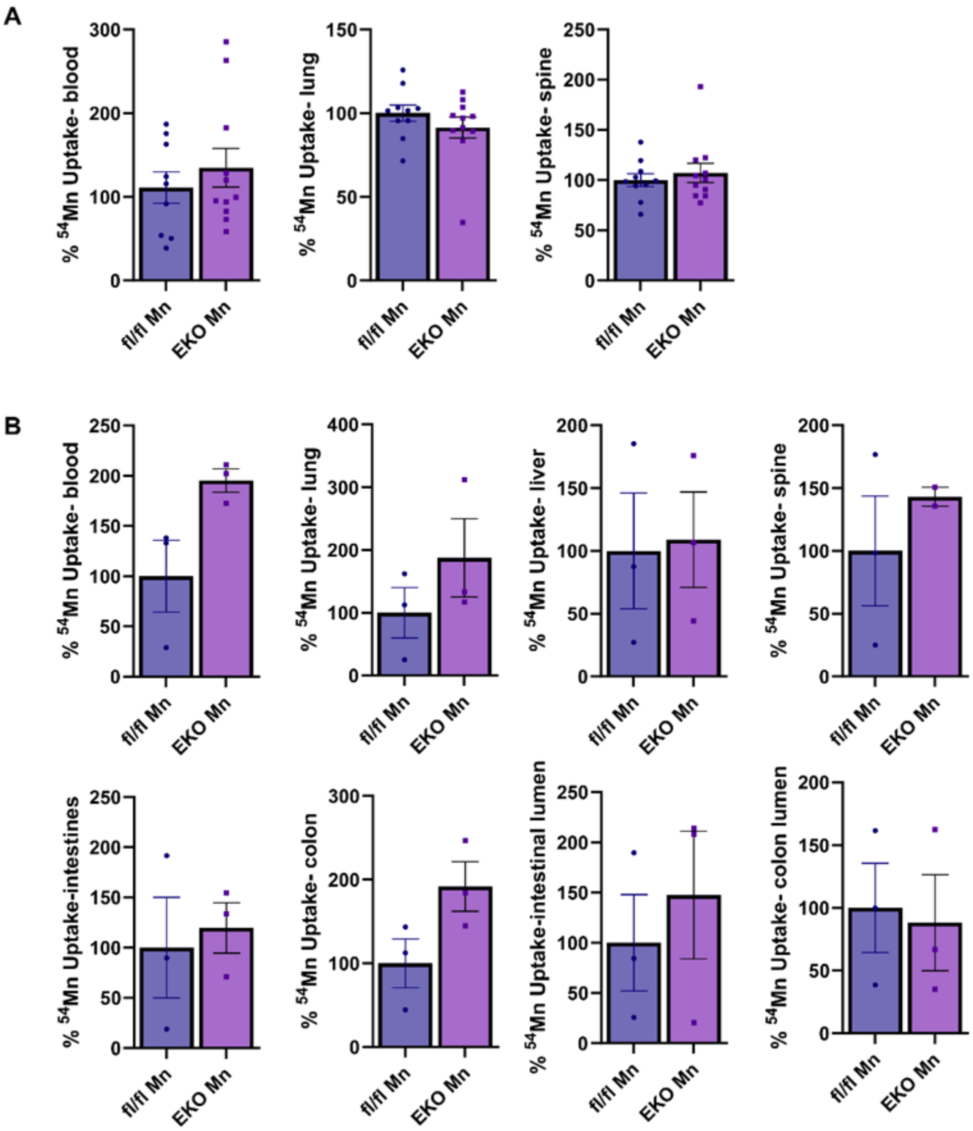


**Supplementary Figure 9.** Tissue %$^{54}$Mn uptake levels after dual delivery following nasal Mn supplementation. fl/fl and EKO mice received daily nasal Mn supplementation for 4 weeks. A) $^{54}$Mn was delivered via nasal delivery, and tissues were collected 1h later. B) $^{54}$Mn was delivered via subcutaneous injections, and tissues were collected 3h later. Radioactivity (counts per minute, cpm) was measured using a gamma counter and normalized to tissue weight.

EKO cpm values are presented as a percentage relative to fl/fl cpm. Data are presented as mean ± SEM; statistical significance was determined using Student's t-test.

Supplementary Figure 10. ZIP14 expression in hCMEC/D3 cells.

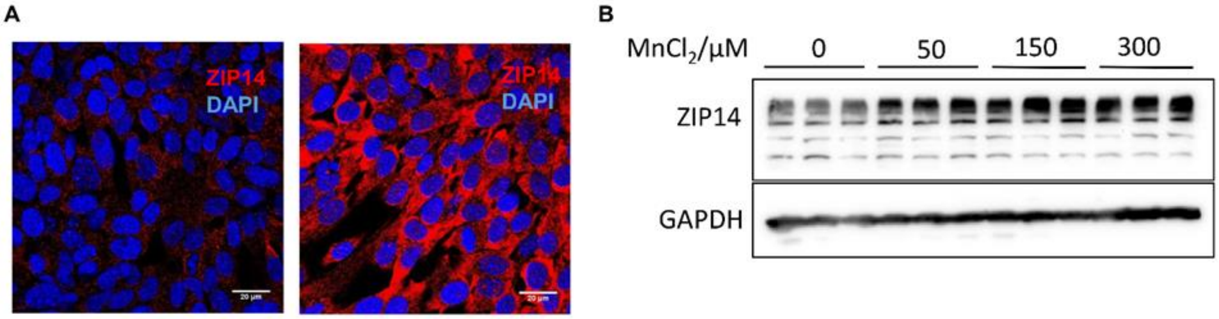


**Supplementary Figure 10.** ZIP14 expression in hCMEC/D3 cells. A) Representative immunofluorescence images of ZIP14 (red) staining in wild-type hCMEC/D3 cells without Mn supplementation (left) and after 72h Mn supplementation (right). B) Western blot showing ZIP14 protein expression in wild-type hCMEC/D3 cells treated with increasing concentrations of $MnCl_2$. GAPDH serves as the loading control. n=3 for each concentration.

Supplementary Figure 11. Diagram of potential routes of nasal delivery

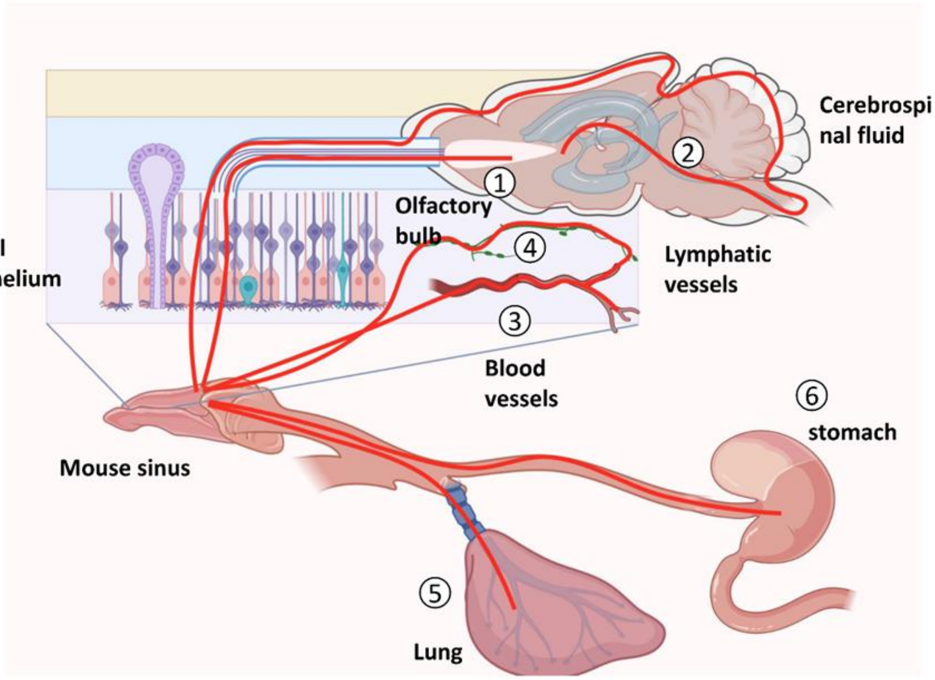


**Supplementary Figure 11.** Diagram of potential routes of nasal delivery. After entering the nasal cavity, substances may: 1) be taken up by olfactory sensory neurons and through to the olfactory bulb, to the central nervous system; or 2) travel paracellularly between olfactory epithelial cells into the cerebrospinal fluid of the subarachnoid space, where it can travel to the ventricles and diffuse into the central nervous system; or leak into the 3) blood vessels or 4) lymphatic vessels surrounding the nasal epithelium; or 5) be inhaled into the lungs; or 6) leak into the stomach.
